# Neuronal mechanism of a BK channelopathy in absence epilepsy and movement disorders

**DOI:** 10.1101/2021.06.30.450615

**Authors:** Ping Dong, Yang Zhang, Mohamad A. Mikati, Jianmin Cui, Huanghe Yang

**Author notes:** Corresponding author: Huanghe Yang.

## Abstract

A growing number of gain-of-function (GOF) BK channelopathy have been identified in patients with epilepsy and paroxysmal movement disorders. Nevertheless, the underlying pathophysiology and corresponding therapeutics remain obscure. Here we utilized a knock-in mouse model carrying human BK-D434G channelopathy to investigate the neuronal mechanism of BK GOF in the pathogenesis of epilepsy and movement disorders. We found that the BK-D434G mice manifest the clinical features of absence epilepsy and exhibit severe motor deficits. BK-D434G mutation causes hyperexcitability of cortical pyramidal neurons and cerebellar Purkinje cells, which contributes to the pathogenesis of absence seizures and the motor defects, respectively. A BK channel blocker paxilline potently suppresses BK-D434G-induced hyperexcitability and effectively mitigates absence seizures in mice. Our study thus uncovered a neuronal mechanism of BK GOF in absence epilepsy and provided the evidence that BK inhibition is a promising therapeutic strategy to mitigate BK GOF-induced neurological disorders.

**Significance:** Dysfunction of BK channels or BK channelopathy has been increasingly implicated in diverse neurological disorders including epilepsy, movement, cognitive and neurodevelopmental disorders. However, precision medicine to treat BK channelopathy is lacking. Here we characterized a mouse model carrying a gain-of-function BK channelopathy D434G from a large family of patients with absence epilepsy and involuntary movement disorders. The BK-D434G mice resemble the clinical manifestations of absence seizures and exhibit severe motor defects. The hyperexcitability in BK-D434G cortical neurons and cerebellar Purkinje cells underscores the neuronal mechanism of BK gain-of-function induced absence epilepsy and movement disorders. The effectiveness of a BK channel blocker on preventing absence seizures suggests that BK inhibition is a promising strategy to treat gain-of-function BK channelopathy.

## Introduction

*KCNMA1* encodes the pore forming α subunit of the Ca^2+^- and voltage-activated large-conductance BK type potassium channels that are widely expressed in the brain with high expression levels in the cortex, cerebellar Purkinje cells, thalamus, hippocampus, basal ganglia, habenula, and olfactory bulb (1-4). Owing to its large single channel conductance, its dual sensitivity to both voltage and intracellular Ca^2+^ and its spatial proximity to voltage-gated Ca^2+^ channels (VGCCs) (4-9) (Fig. 1A), BK channels play pivotal roles in shaping action potential repolarization, giving rise to fast after-hyperpolarization (fAHP), controlling dendritic Ca^2+^ spikes and influencing synaptic transmission (1, 2, 10-12). Consistent with its importance in the nervous system, dysfunction of BK channels has been implicated in the pathophysiology of various neurological disorders including epilepsy (12-16), movement disorders (13, 15, 17-22), and neurodevelopmental and cognitive disorders such as intellectual delay (15, 16, 18, 21, 23), autism spectrum disorder (14, 17, 21, 24), Fragile X syndrome (25) and Angelman syndrome (26). How BK channels involve in such a diverse spectrum of neurological disorders (27-29), however, remains largely elusive and demands in-depth studies.

**Fig. 1.**
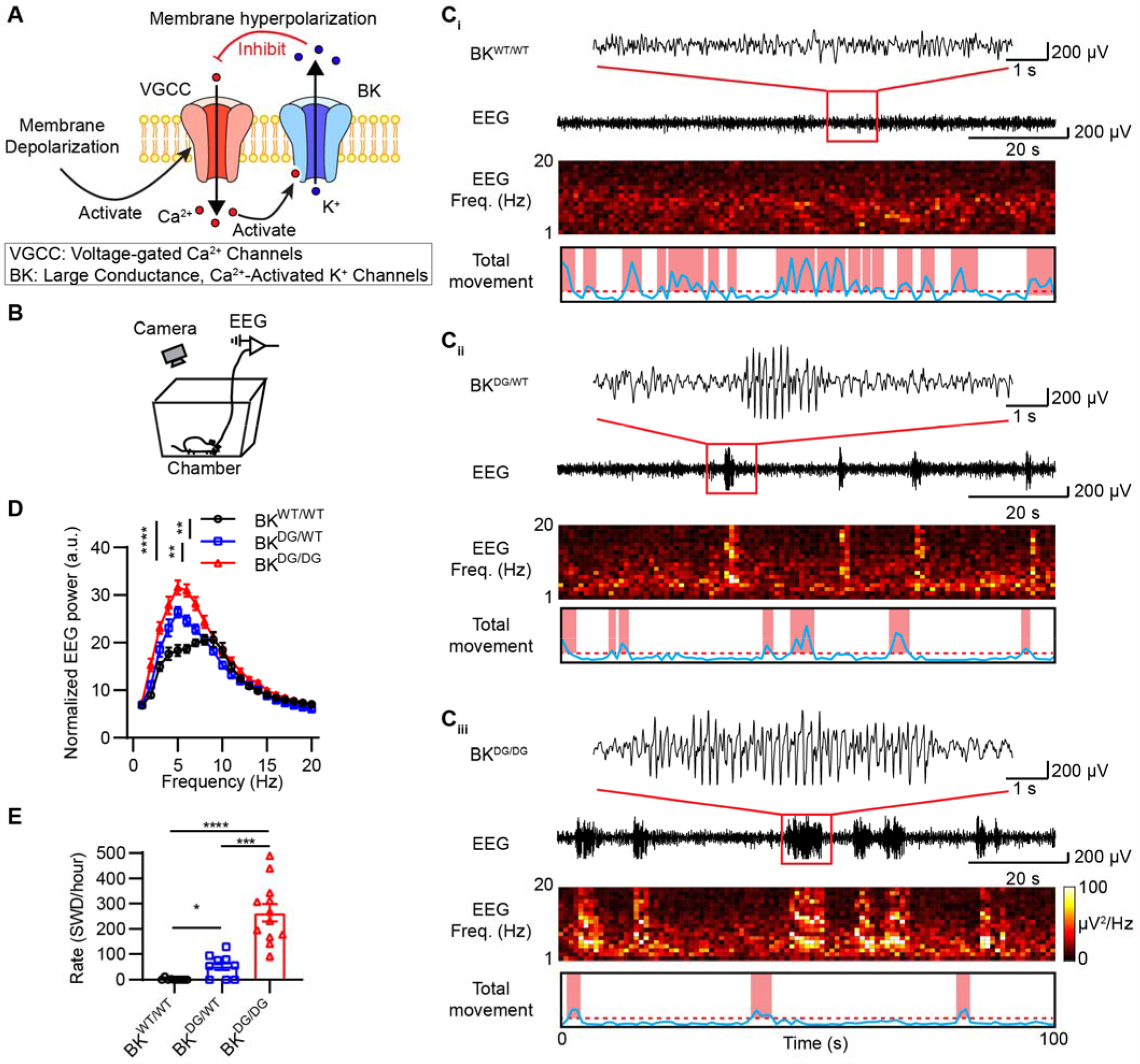
BK-D434G knock-in mice have spontaneous absence seizures. (**A**) Ca^2+^- and voltage-activated BK channels control the opening of voltage-gated Ca^2+^ channels (VGCCs or Ca_V_s) through a negative-feedback mechanism. (**B**) Schematic of the simultaneous video-electroencephalogram (EEG) recordings of freely moving mice. (**C**) *BK*^*DG/WT*^ (**C**_**ii**_) and *BK*^*DG/DG*^ (**C**_**iii**_) but not *BK*^*WT/WT*^ mice (**C**_**i**_) had spontaneous spike-wave discharges (SWDs) and frequent behavior arrest. Top: raw EEG traces. Red rectangles show the corresponding EEG traces on an expanded time scale. Middle: corresponding spectrograms of the EEG traces. Bottom: video-based analysis of the total movement. The behavior status is classified as motion state (red boxes) or immobile state (white boxes). See Methods for details. (**D**) Summary of power spectral density of EEG recorded from *BK*^*WT/WT*^ (n = 9), *BK*^*DG/WT*^ (n = 9) and *BK*^*DG/DG*^ (n = 12) mice. Normalization was performed by averaging the power to the total recording time. Two-way ANOVA, *F*_(2,27)_ = 9.683, *P* = 0.0007. (**E**) Summary of the number of spontaneous SWDs per hour for from *BK*^*WT/WT*^ (n = 9), *BK*^*DG/WT*^ (n = 9) and *BK*^*DG/DG*^ (n = 12) mice. One-way ANOVA test, *F*_(2,27)_ = 11.57, *P* = 0.0002. * P < 0.05, \*\*\**P* < 0.001, \*\*\*\**P* < 0.0001. In all plots and statistical tests, summary graphs show mean ± s.e.m.

*KCNMA1* variants identified from human genetic analysis provide unique opportunities to understand the neurological functions of BK channels (27, 29). The first *KCNMA1*-linked potassium channelopathy D434G from a large family of patients with generalized epilepsy and/or paroxysmal dyskinesia (13). Of the sixteen BK-D434G patients, nine individuals had absence epilepsy, which is characterized by sudden, brief lapses of consciousness accompanied by behavioral arrest and distinctive bilaterally synchronous spike-and-wave discharges (SWDs) at 2.5-4 Hz (30); twelve BK-D434G patients developed paroxysmal nonkinesigenic dyskinesia (PNKD), which is an episodic movement disorder characterized by attacks of hyperkinesia with intact consciousness (13). Interestingly, five BK-D434G patients were affected by both absence epilepsy and PNKD. Subsequent biophysical characterizations demonstrated that BK-D434G is a gain-of-function (GOF) mutation with enhanced Ca^2+^ sensitivity (13, 31-33). It is intriguing why a GOF potassium channel mutation is associated with epilepsy and dyskinesia, which is characterized by hyperexcitability and hypersynchronization in nature. A growing number of human *KCNMA1* variants have been identified over the past several years (27, 29). However, it is unknown 1) whether the *KCNMA1* variants cause the associated neurological disorders; 2) how the *KCNMA1* variants affect neuronal activities at cellular level; 3) whether targeting the mutant BK channels is effective to mitigate the associated neurological symptoms.

To address these questions, we characterized a knock-in mouse model carrying the BK-D434G mutation. We found that the BK-D434G mice align with the clinical manifestations of absence seizures and response to anti-absence medications. *In vitro* brain slice recordings revealed that the hyperexcitability of the cortical pyramidal neurons contribute to BK-D434G-induced spontaneous absence seizures. The effectiveness of paxilline (PAX), a BK channel specific blocker, on suppressing BK-D434G-induced absence seizures in mice establishes the causal relationship of this GOF and absence seizures. BK-D434G also induces hyperactivity in Purkinje cells and leads to functional and morphological changes, all of which contribute to the observed motor defects. Our study not only elucidate the cellular basis of the BK-D434G channelopathy in epilepsy and movement disorders, but also demonstrate that BK inhibition can be a promising therapeutic strategy to mitigate BK GOF-induced epilepsy.

## Results

### BK-D434G knock-in mice resemble clinical manifestations of absence epilepsy

A knock-in mouse line carrying BK-D434G mutation was generated by homologous recombination (Fig. S1*A* and Methods). The resulting animals were confirmed by both genotyping PCR (Fig. S1*B*) and genomic sequencing (Fig. S1*C*). The heterozygous BK-D434G mutation (*BK*^*DG/WT*^) mice were viable and survived into adulthood. However, only 17.3% of the offspring were homozygous *BK*^*DG/DG*^ under the *BK*^*DG/WT*^ × *BK*^*DG/WT*^ breeding scheme, which was significantly less than the expected 25% Mendelian inheritance (*P* < 0.05, Chi-square = 6.495, Fig. S1*D*). This indicates that the BK-D434G mutation homozygosity had some detrimental effects to the *BK*^*DG/DG*^ mice.

Nine out of sixteen of the BK-D434G channelopathy patients had generalized epilepsy (13). These patients typically had absence seizures with spike-wave-discharges (SWDs). We therefore used simultaneous video-electroencephalogram (EEG) recording (Fig. 1*B* and Methods) to examine if the knockin mice resemble the human BK-D434G patients’ clinical manifestations. We found that both the heterozygous *BK*^*DG/WT*^ and homozygous *BK*^*DG/DG*^ mice, but not the *BK*^*WT/WT*^ control mice exhibited frequent episodes of spontaneous, generalized SWDs, each of which lasted for 0.5-10 seconds (Fig. 1*C*, Table 1 and Movie S1). Power spectral analysis of the SWDs showed that the epileptic events of the BK-D434G mice were composed of strong frequency bands of 3-8 Hz (Fig. 1*D*), which is comparable to the typical SWD frequency range in other rodent models with absence seizures (Table 1) (34, 35). Compared with the *BK*^*DG/WT*^ mice, which had 54.1 ± 13.4 SWDs per hour, the homozygous *BK*^*DG/DG*^ mice showed dramatically increased incidences of SWDs (263.7 ± 34.8 SWDs/hour) (Fig. 1*E*). This suggests that BK-D434G homozygosity can lead to more severe phenotypes, which may contribute to the increased lethality of the *BK*^*DG/DG*^ mice (Fig. S1*D*).

**Table 1.**
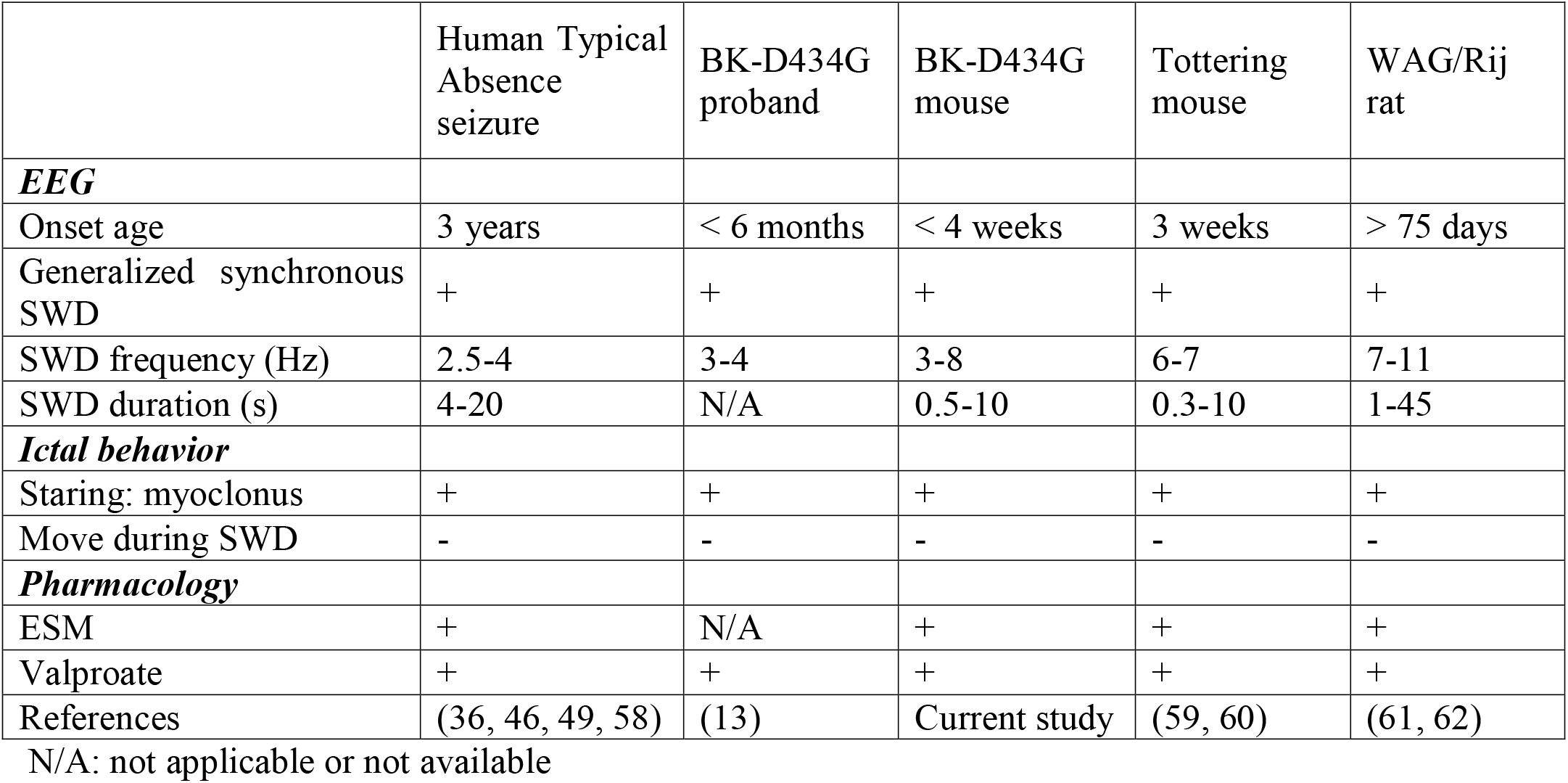
Comparison of the absence seizures in rodent models and human patients.

|  | Human Typical<br>Absence<br>seizure | BK-D434G<br>proband | BK-D434G<br>mouse | Tottering<br>mouse | WAG/Rij<br>rat |
| --- | --- | --- | --- | --- | --- |
| ***EEG*** |  |  |  |  |  |
| Onset age | 3 years | < 6 months | < 4 weeks | 3 weeks | > 75 days |
| Generalized synchronous<br>SWD | + | + | + | + | + |
| SWD frequency (Hz) | 2.5-4 | 3-4 | 3-8 | 6-7 | 7-11 |
| SWD duration (s) | 4-20 | N/A | 0.5-10 | 0.3-10 | 1-45 |
| ***Ictal behavior*** |  |  |  |  |  |
| Staring: myoclonus | + | + | + | + | + |
| Move during SWD | - | - | - | - | - |
| ***Pharmacology*** |  |  |  |  |  |
| ESM | + | N/A | + | + | + |
| Valproate | + | + | + | + | + |
| References | (36, 46, 49, 58) | (13) | Current study | (59, 60) | (61, 62) |

N/A: not applicable or not available

Combining the EEG recording with an unbiased, automatic video analysis on the total movement on the freely moving mice (Fig. S2 and Methods), we demonstrated that the BK-D434G mice exhibited frequent behavioral arrest during the SWDs onset (Movie S1 and Table 1), another hallmark of absence epilepsy (36). Different from the *BK*^*WT/WT*^ mice that constantly underwent alternating locomotive and non-locomotive status (Fig. 1*C*_*i*_), the *BK*^*DG/WT*^ and *BK*^*DG/DG*^ mice showed much higher incidences and longer durations of non-locomotion (Fig. 1*C*_*ii*_ and 1*C*_*iii*_). After aligning the mouse locomotive activities with the SWDs, we found that when SWDs developed, the mice were behaviorally arrested; whereas when the mice were spared from SWDs, they were able to freely move around.

As the BK-D434G proband responded to valproate (13), we tested the effects of the first-line anti-absence medicines valproate and ethosuximide (ESM) on our *BK*^*DG/DG*^ mice. Administration of valproate or ESM effectively suppressed the frequent SWDs in the animals for about an hour (Fig. 2). Typical SWDs accompanied by behavioral arrest and responsiveness to the first-line anti-absence seizure medicines (Table 1) explicitly demonstrated that the BK-D434G mice fully align with the clinical manifestations of absence epilepsy from the human patients carrying the BK-D434G mutation (13).

**Fig. 2.**
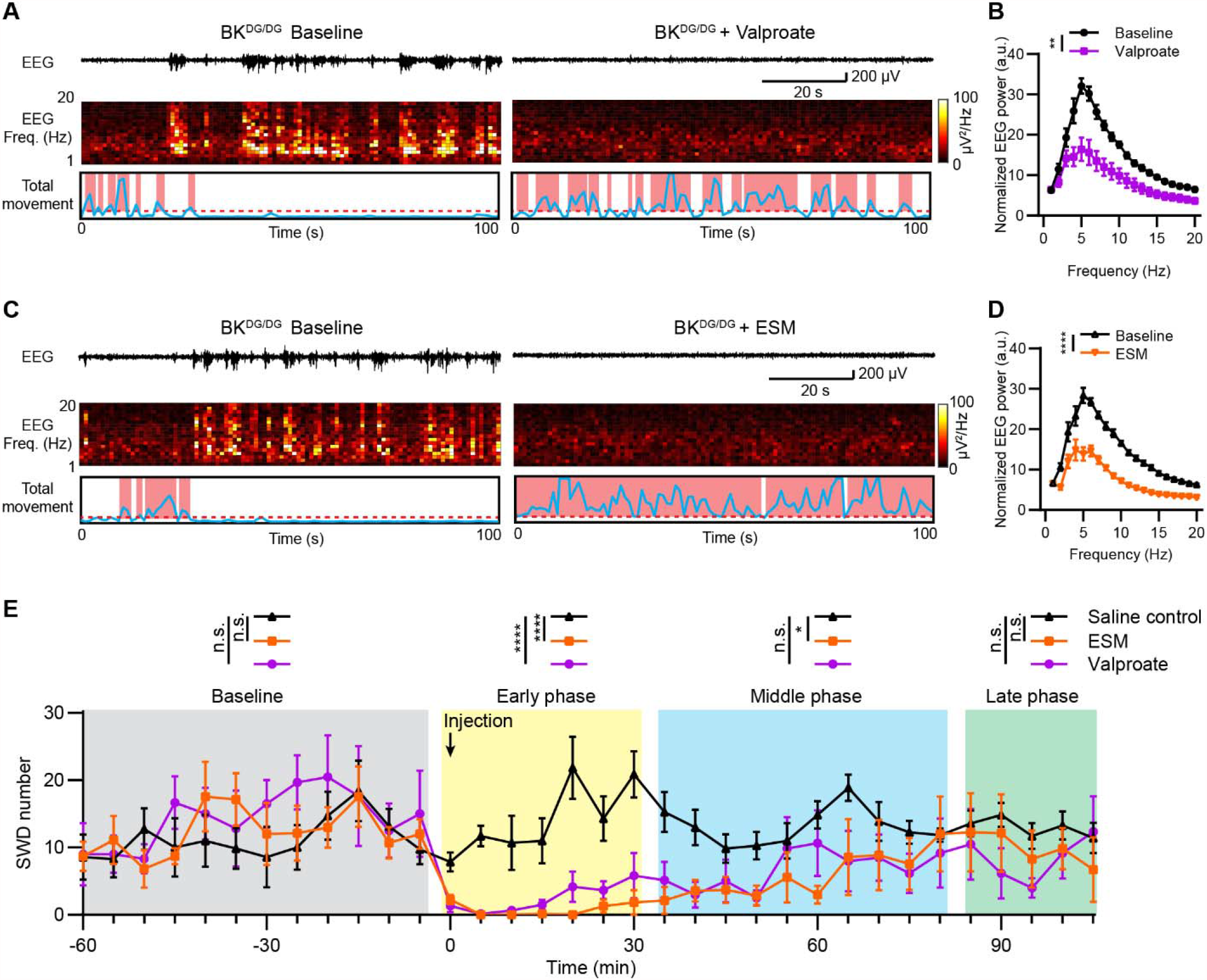
First-line anti-absence seizure medicines can abolish the absence seizure in BK-D434G mouse. (**A, C**) Representative EEG traces, corresponding spectrograms, and total movement from the *BK*^*DG/DG*^ mice before and after the first-line anti-absence seizure medicines valproate (**A**) or ESM (**C**) administration. (**B, D**) Summary of power spectral density of EEG recorded from *BK*^*DG/DG*^ mice during before and after valproate (**B**, 200 mg/kg, Two-way ANOVA, *F*_(1,10)_ = 13.30, *P* = 0.0045. n = 6 mice) or ESM (**D**, 150 mg/kg, Two-way ANOVA, *F*_(1,12)_ = 66.10, *P* < 0.0001. n = 7 mice) administration. (**E**) Time course of the drug effects of ethosuximide (ESM, orange), valproate (purple) and saline control (black) on the spontaneous SWDs of the *BK*^*DG/DG*^ mice. (Bin size = 5 min). The drug effects were empirically divided into 4 different phases: baseline phase, 60 min prior to injection (grey box, two-way repeated-measures ANOVA, F_(2,17)_ = 1.176, *P* = 0.3324); early phase, 30 min post injection (yellow box, two-way repeated-measures ANOVA, F_(2,17)_ = 78.61, *P* < 0.0001); middle phase, from 35 to 80 min post injection (blue box, two-way repeated-measures ANOVA, F_(2,17)_ = 3.906, *P* = 0.0402); and late phase, from 85 to 105 min post injection (green box, two-way repeated-measures ANOVA, F_(2,17)_ = 0.5274, *P* = 0.5274). n = 7 mice per group for saline control, ESM, n = 6 mice for valproate administration. The error bars indicate s.e.m., * *P* < 0.05, ** *P* < 0.01, *** *P* < 0.001, **** *P* < 0.0001.

### BK-D434G knock-in mice are susceptible to convulsant-induced tonic-clonic seizures

In addition to absence seizures, two BK-D434G channelopathy patients were also reported to develop generalized tonic-clonic seizures (13). We hypothesized that BK-D434G GOF mutation may increase the susceptibility to develop tonic-clonic seizures. To test this, we administered pentylenetetrazole (PTZ), a convulsant (37), to the *BK*^*WT/WT*^ and *BK*^*DG/WT*^ mice. We found that the low dosage of PTZ injection (40 mg/kg) induced generalized seizure (GS) stage (seizure score ≥ 4, see Methods for details) in all *BK*^*DG/WT*^ mice, whereas the same dosage of PTZ injection induced GS stage in only 42.9% of the *BK*^*WT/WT*^ mice (Fig. S3*A*). Furthermore, the *BK*^*DG/WT*^ mice exhibited significantly increased seizure scores (Fig. S3*A*), markedly prolonged GS duration (Fig. S3*B*) and dramatically reduced latency to GS (Fig. S3*C*). Our characterizations indicated that the BK-D434G mice not only have spontaneous seizures, but also are more vulnerable to PTZ-induced generalized tonic-clonic seizures.

### Cortical pyramidal neurons of BK^DG/WT^ mice show hyperexcitability

Cortical neurons play essential roles in the pathogenesis of absence seizures (30); and BK channels are highly expressed in cortical pyramidal neurons (2, 3). Therefore, we investigated whether BK-D434G GOF mutation alters the membrane excitability of cortical pyramidal neurons. Our acute brain slice recording showed that the cortical pyramidal neurons from the *BK*^*DG/WT*^ mice exhibited hyperexcitability as evidenced by the significantly increased action potential frequency compared with the *BK*^*WT/WT*^ mice (Fig. 3, *A-B* and Table S1). Single action potential analysis of the first spikes revealed that the *BK*^*DG/WT*^ cortical neurons exhibited much faster repolarization as evidenced by the significantly shortened action potential duration (AP90) and augmented after-hyperpolarization amplitude (AHP) (Fig. 3, *C-E*). As K^+^ efflux through BK channels contributes to fast after-hyperpolarization (fAHP) (11, 12), our observations of steeper repolarization and enhanced AHP in the *BK*^*DG/WT*^ neurons suggest that the BK-D434G GOF mutant channels, which have a higher Ca^2+^ sensitivity (13, 31), can more efficiently hyperpolarize the membrane following membrane depolarization and VGCC opening. The faster and stronger hyperpolarization induced by BK-D434G would enable faster recovery of the voltage-gated sodium channels (Na_V_) (Fig. 3*F*) and potentially facilitate the activation of the hyperpolarization-activated cation (HCN) channels (11, 38). The rapid repriming of the Na_V_ channels and enhanced HCN channel activation collectively enables the cortical neurons to fire at a higher frequency. Taken together, our electrophysiological characterization of the cortical pyramidal neurons from the BK-D434G mice demonstrated a neuronal mechanism of BK GOF-induced hyperexcitability.

**Fig. 3.**
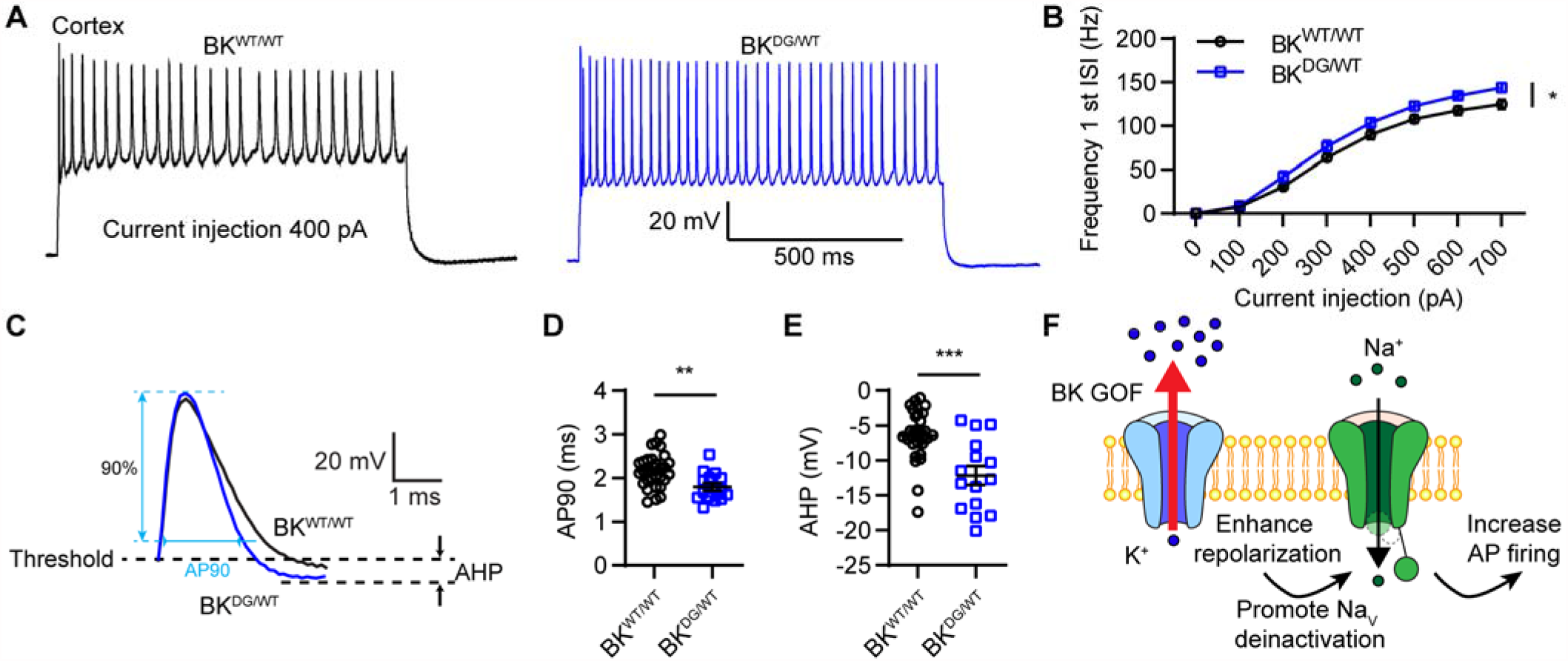
Whole-cell electrophysiology shows that the BK-D434G cortical pyramidal neurons are hyperactive. (**A**) Representative evoked action potentials in cortical pyramidal neurons from *BK*^*WT/WT*^ and *BK*^*DG/WT*^ mice. Firing was elicited by a 400 pA current injection for 1s. (**B**) Action potential frequency of the first inter-spike interval (1^st^ ISI) from the *BK*^*WT/WT*^ and *BK*^*DG/WT*^ cortical neurons. Two-way repeated-measures ANOVA, *F*_(1,42)_ = 6.640, *P* = 0.0136. (**C**) Representative single action potential waveforms elicited by 400 pA current injection. Definitions of action potential parameters are labeled with cyan and black dash line. AP90 was used to define as action potential duration of 90% repolarization and AHP denotes after hyperpolarization. (**D, E**) *BK*^*DG/WT*^ cortical neurons have shorter action potential duration (**D**, two-tailed unpaired Student’s *t*-test, t_42_=3.111, *P* = 0.0033) and higher amplitude of fast AHP (**E**, two-tailed unpaired Student’s *t*-test, t_42_=4.294, *P* = 0.0001) compared with *BK*^*WT/WT*^ neurons. n = 29 neurons from five *BK*^*WT/WT*^ mice and n = 15 neurons from four *BK*^*DG/WT*^ mice. (**F**) The gain-of-function BK-D434G mutant channels enhance membrane repolarization and accelerate de-inactivation of the voltage-gated sodium (Na_V_) channels, which enable the excitatory neurons to fire at a higher frequency. * *P* < 0.05, ** *P* < 0.01, *** *P* < 0.001. In all plots and statistical tests, summary graphs show mean ± s.e.m.

### Pharmacological inhibition of BK channels suppresses BK-D434G-induced seizures

We next tested whether pharmacological inhibition of BK channels can restore normal firing and suppress absence seizures in the BK-D434G mice. PAX, a BK channels specific blocker, effectively suppressed the hyperexcitability of cortical pyramidal neuron (Fig. 4, *A* and *B* and Table S1), markedly slowed down membrane repolarization, prolonged AP90 and reduced AHP amplitude (Fig. 4, *C-E*). Consistent with our brain slice recording, administration of PAX (0.35 mg/kg i.p.) eliminated the spontaneous SWDs of the *BK*^*DG/DG*^ mice and prevented their behavioral arrest for about 30 min (Fig. 4, *F-H*). Moreover, we found the PAX also decreased the severity of the PTZ-induced seizures in the BK-D434G heterogeneous mice (Fig. S4). Compared with the saline control, PAX administration significantly decreased seizure scores (Fig. S4*A*), markedly reduced GS duration (Fig. S4*B*) and dramatically prolonged the latency to GS (Fig. S4*C*). All these are consistent with the anti-convulsant effect of PAX on the rodent models of epilepsy, including the PTZ-injected rodent models and an Angelman syndrome mouse model with enhanced BK channel activity (39-41). Ou*r in vitro* and *in vivo* experiments thus explicitly showed that pharmacological inhibition of BK channels can suppress absence seizures in the BK-D434G mice and reduce their vulnerability to convulsant-induced seizure. Our findings not only further supported the causal effect of BK-D434G GOF in epilepsy, but also demonstrated pharmacological inhibition as a promising therapeutic strategy to mitigate BK GOF induced epilepsy.

**Fig. 4.**
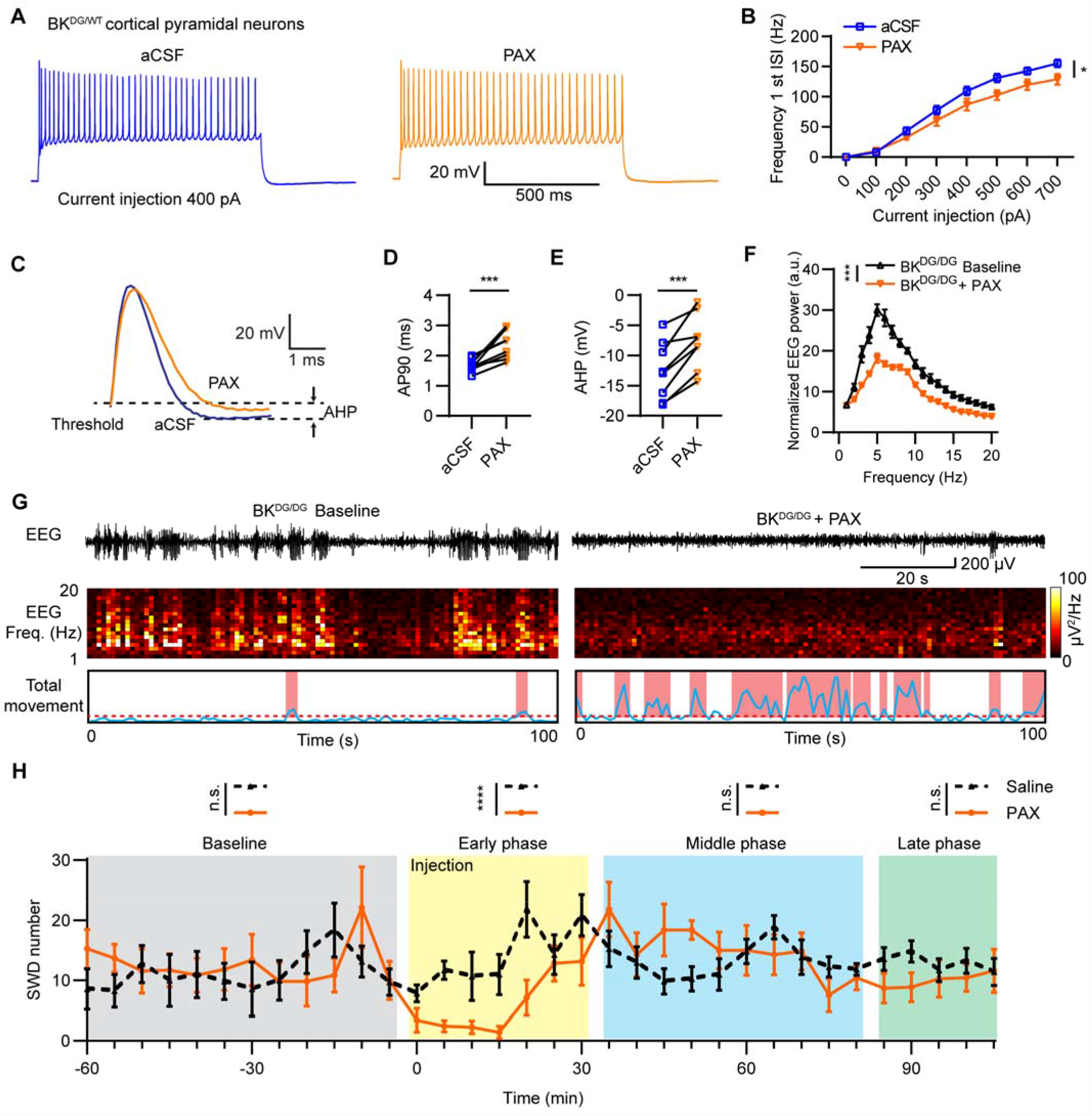
Paxilline (PAX), a BK channel blocker, reduces the hyperactivity of the cortical pyramidal neurons and suppresses the spontaneous absence seizures of the BK-D434G mice. (**A**) Representative evoked action potentials in cortical pyramidal neurons from *BK*^*DG/WT*^ mice. Firing was elicited by 1s, 400 pA current injection, before and after application of 10 µM PAX. (**B**) Action potential frequency of the first inter-spike interval (1^st^ ISI) for the *BK*^*DG/WT*^ cortical neurons before and after application of PAX. Two-way repeated-measures ANOVA, *F*_(1,14)_ = 4.799, *P* = 0.0459. n = 8 neurons from 3 mice. (**C**) Representative single action potential waveforms of the *BK*^*DG/WT*^ cortical neurons elicited by 400 pA current injection before and after application of PAX. (**D, E**) PAX broadens action potential (AP) duration (**D**, two-tailed paired Student’s *t*-test, t_7_=4.591, *P* = 0.0025) and suppresses after-hyperpolarization (AHP) in the cortical pyramidal cells from BK^DG/WT^ mice (**E**, two-tailed paired Student’s *t*-test, t_7_=5.581, *P* = 0.0008). n = 8 neurons from 3 mice. (**F**) Summary of power spectral density of EEG recorded from *BK*^*DG/DG*^ mice before and after PAX administration (n = 7 mice). Two-way ANOVA, *F*_(1,12)_ = 26.39, *P* = 0.0002. n = 7 mice. (**G**) Representative EEG traces, corresponding spectrograms, and total movement from the *BK*^*DG/DG*^ mice before (left panel) and after 0.35 mg/kg PAX (right panel) administration. (**H**) Time course of the drug effects of PAX (orange line) and saline control (black dash line) on the spontaneous SWDs of the *BK*^*DG/DG*^ mice. (Bin size = 5 min). PAX is effective in the early phase, 30 min post injection (yellow box, two-way repeated-measures ANOVA, F_(1,12)_ = 42.57, *P* < 0.0001), but not in the later phases. n = 7 mice per group. The error bars indicate s.e.m., * *P* < 0.05, *** *P* < 0.001, **** *P* < 0.0001. In all plots and statistical tests, summary graphs show mean ± s.e.m.

### BK-D434G mice exhibit severe locomotive defects

In addition to absence epilepsy, majority of the patients (twelve out of sixteen) with BK-D434G mutation also had paroxysmal movement dyskinesia (13). We thus performed a battery of locomotor tests to assess the potential motor defects of the knock-in mice. We first used the open field test to evaluate their general locomotor activities (Fig. 5*A*). During a 15-minute test, the total travel distance of the *BK*^*DG/DG*^ mice were dramatically less than that of the *BK*^*WT/WT*^ and *BK*^*DG/WT*^ mice (Fig. 5 *A* and *B*). Our balance beam test (Fig. 5*C* and Movie S2) demonstrated that the BK^*DG/DG*^ mice took significantly longer time to traverse the balance beam (Fig. 5*D*) and had significant more incidences of hind-limb slips compared with the WT controls (Fig. 5*E*). The *BK*^*DG/WT*^ mice showed no defect on transverse time yet had a milder defect on the number of hind-limb slips. Interestingly, the *BK*^*DG/WT*^ mice occasionally but the *BK*^*DG/DG*^ mice always used their tails to maintain their balance on the beam (Fig. 5*C* and Movie S2). We next performed accelerated rotarod test, which is a standard assay to evaluate impairment in rodent motor performance. Both the BK^*DG/WT*^ and the BK^*DG/DG*^ mice performed poorly on this more challenging motor task with significantly shorter latency to fall (Fig. 5*F* and Movie S3). Compared with the *BK*^*DG/WT*^ mice, the *BK*^*DG/DG*^ mice showed worst performance on rotarod. The severe defects observed during the accelerated rotarod test clearly showed that the BK-D434G mice indeed have impaired motor functions. Several factors such as muscle strength, motor learning and motor coordination may affect rotarod performance (42). To specifically evaluate the motor coordination functions of the BK-D434G mice, we performed gait analysis utilizing footprints (Fig. 5*G*). Our result revealed that the BK^*DG/DG*^ mice had significantly shorter hind-limb stride lengths than the BK^*WT/WT*^ mice (Fig. 5*H*), suggesting that these mutant animals had severe defects on motor coordination. Taken together, our multiple locomotor tests clearly demonstrated that the motor functions of the BK-D434G mice were severely impaired.

**Fig. 5.**
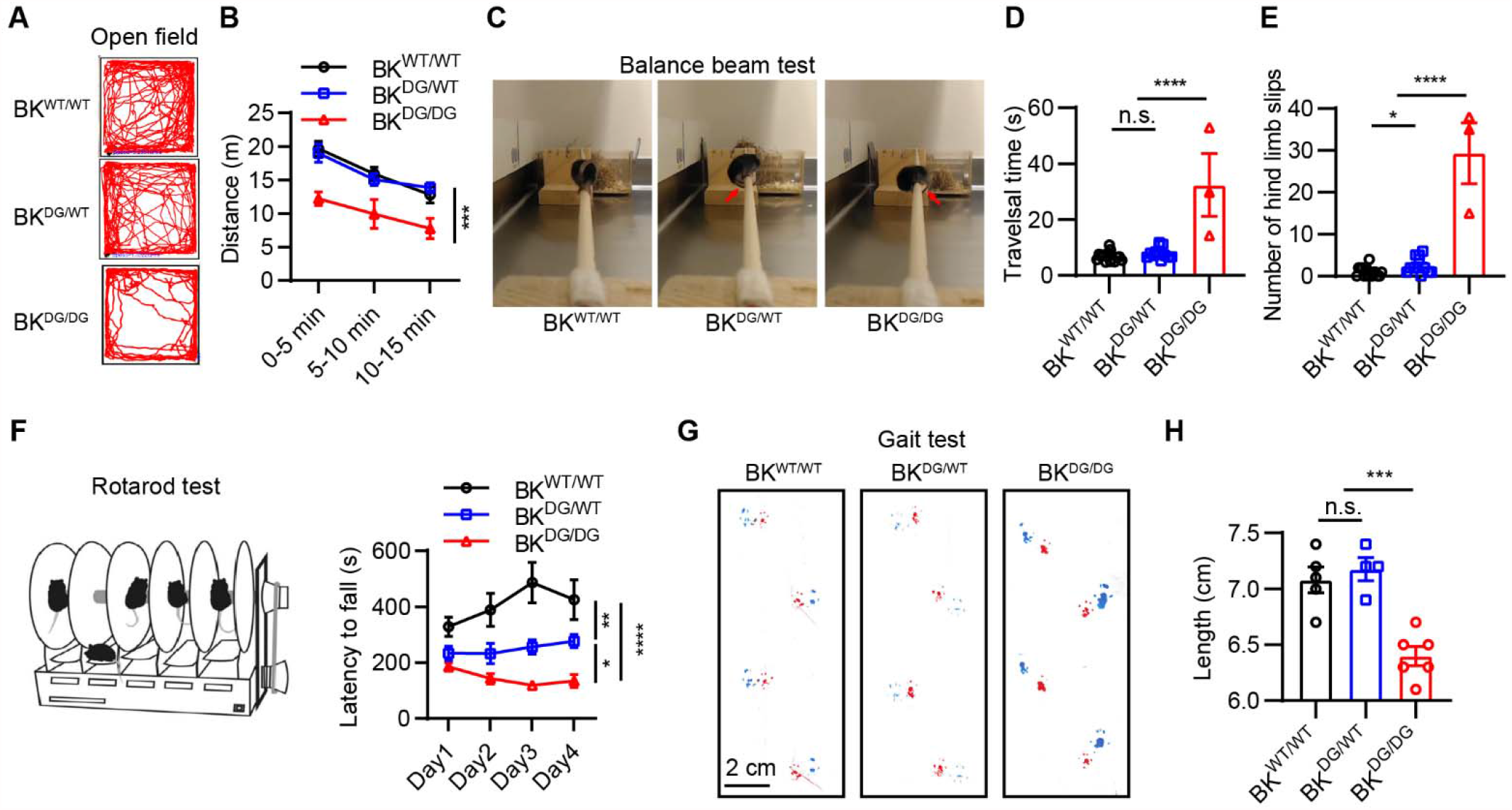
BK-D434G mice exhibit motor defects. (**A**) Representative animal track in the open-field chamber of BK^WT/WT^, BK^DG/WT^ and BK^DG/DG^ mice. (**B**) BK^DG/DG^ mice showed decreased locomotor activity in 15 min open field test. Two-way repeated-measures ANOVA, *F*_(2,23)_ = 9.974, *P* = 0.0008. BK^WT/WT^, n = 9 mice, BK^DG/WT^, n = 12 mice, BK^DG/DG^, n = 5 mice. (**C**) Balance beam test of BK-D434G mutation mice, the red arrows indicate the low position of the tails when hind-limb slips caused balance loss were observed during the test. (**D, E**) BK^DG/DG^ mice had significantly more hind-limb slips on the balance beams (**D**, One-way ANOVA test, *F*_(2,27)_ = 75.05, *P* < 0.0001) and took significantly longer to traverse the balance beam (**E**, One-way ANOVA test, *F*_(2,27)_ = 27.14, *P* < 0.0001) compared with BK^WT/WT^ controls. BK^WT/WT^, n = 15 mice, BK^DG/WT^, n = 12 mice, BK^DG/DG^, n = 3 mice. (**F**) Accelerating rotarod latency to fall times for BK-D434G mutation mice over four days of testing compared with control mice, showing a significant deficit for BK-D434G mice in this test. ****p < 0.0001, significant difference between genotypes for those trials. Two-way repeated-measures ANOVA, *F*_(2,28)_ = 19.56, *P* < 0.0001. BK^WT/WT^, n = 8 mice, BK^DG/WT^, n = 12 mice, BK^DG/DG^, n = 11 mice. (**G**) Representative images of the gait patterns of the BK^WT/WT^, BK^DG/WT^ and BK^DG/DG^ mice, with forepaws are represented by red paint and hind-paws by blue paint (scale bar, 2 cm). (**H**) Quantification reveals shortened stride length. One-way ANOVA test, *F*_(2,12)_ = 18.50, *P* = 0.0002. BK^WT/WT^, n = 5 mice, BK^DG/WT^, n = 4 mice, BK^DG/DG^, n = 6 mice. In all plots and statistical tests, summary graphs show mean ± s.e.m., * p<0.05, ** p<0.01, ***p<0.001, ****p<0.0001.

### Hyperexcitability of BK-D434G cerebellar Purkinje cells contributes to motor defects

BK channels are highly expressed in Purkinje cells (PCs) and play a critical role in controlling PC excitability (2). Genetic ablation of BK channels in murine PCs leads to cerebellar ataxia and impaired motor coordination (43, 44) and some KCNMA1 channelopathy patients showed signs of cerebellar atrophy (15, 17-20). Given all these facts, we set out to examine whether BK-D434G GOF in PC contributes to the observed impairments in motor functions. By immunostaining with the PC marker calbindin, we found that the adult BK-D434G mutant mice showed dramatic changes of their PC morphology (Fig. 6*A*). The size of PC soma and the width of PC primary dendrites were significantly enlarged in both the BK^*DG/WT*^ and the BK^*DG/DG*^ mice compared with the BK^*WT/WT*^ mice (Fig. 6*B* and *C*), indicating signs of PC hypertrophy in the BK-D434G mice.

**Fig. 6.**
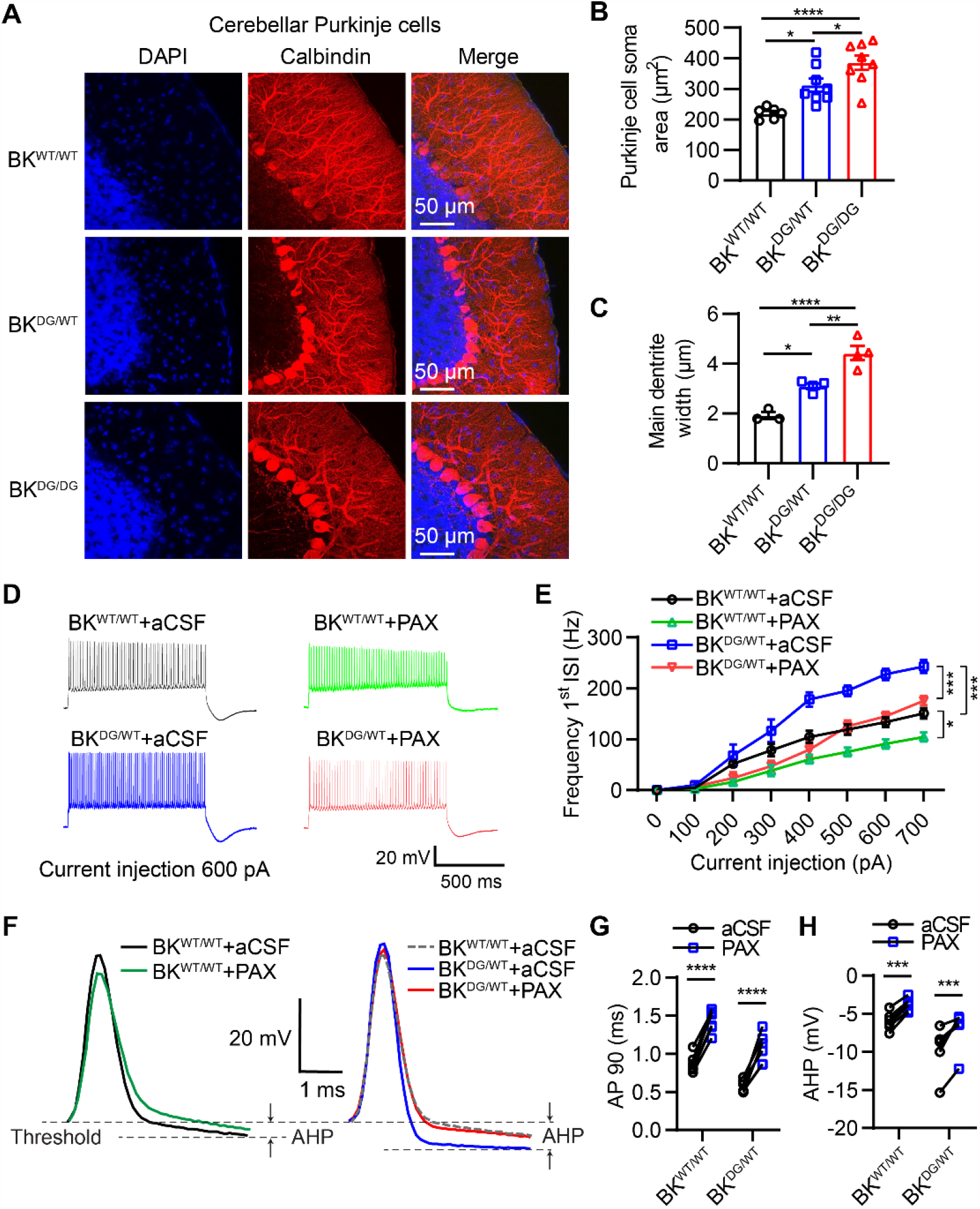
Cerebellar Purkinje cells (PCs) from the BK-D434G mutant mice are hyperactive, which can be suppressed by paxilline. (**A**) Representative confocal images of PCs. The PCs from the BK^DG/WT^ and BK^DG/DG^ mice show signs of hypertrophy. (**B**) The BK^DG/WT^ and BK^DG/DG^ mice have enlarged PC somas. One-way ANOVA test, *F*_(2,19)_ = 14.97, *P* = 0.0001, BK^WT/WT^, n = 6; BK^DG/WT^, n = 8; BK^DG/DG^, n = 8. (**C**) The BK^DG/WT^ and BK^DG/DG^ mice have thickened main dendrite width. The dendrite width is measured at the distance of one cell-soma diameter from the cell soma. One-way ANOVA test, *F*_(2,8)_ = 34.46, *P* = 0.0001, BK^WT/WT^, n = 3; BK^DG/WT^, n = 4; BK^DG/DG^, n = 4. (**D**) Representative evoked action potentials in cerebellar PCs from BK^WT/WT^ and BK^DG/WT^ mice. Firing was elicited by 1s long 600 pA current injection, before and after application of 10 µM PAX. (**E**) Statics of the action potential numbers of BK^WT/WT^ and BK^DG/WT^ PCs, before and after application of PAX. Two-way repeated-measures ANOVA, *F*_(3,20)_ = 20.06, *P* < 0.0001. (**F**) Representative single action potential waveforms elicited by 600 pA current injection for BK^WT/WT^ and BK^DG/WT^ mice before and after 10 µM PAX. In the right panel, BK^WT/WT^ trace is presented in grey dash line for comparison with BK^DG/WT^. (**G, H**) BK^DG/WT^ PCs have shorter action potential duration (AP90) and higher amplitude of fast after-hyperpolarization (fAHP) compared with BK^WT/WT^. PAX broadens AP duration (**G**, Two-way ANOVA, *F*_(1,10)_ = 20.27, *P* < 0.0001) and reduces fAHP (**H**,, Two-way ANOVA, *F*_(1,10)_ = 75.31, *P* < 0.0001) of PCs from both BK^WT/WT^ and BK^DG/WT^ mice. n = 6 neurons from 3 mice per group. In all plots and statistical tests, summary graphs show mean ± s.e.m.. * p<0.05, ** p<0.01, ***p<0.001, ****p<0.0001.

Next, we conducted brain slice patch clamp recording on the PCs from the BK^*WT/WT*^ and the BK^*DG/WT*^ mice (Fig. 6*D-H*). Similar to what we observed in cortical pyramidal neurons (Fig. 3), we found that the BK^*DG/WT*^ PCs had dramatically enhanced firing rate compared with the PCs of the BK^*WT/WT*^ mice (Fig. 6*D* and *E*). The subsequent single action potential waveform analysis showed that the BK^*DG/WT*^ PCs showed faster membrane repolarization (Fig. 6F, right) with significant reduction of action potential duration (Fig. 6*G*) and increase of AHP amplitude (Fig. 6*H*). Consistent with our observations in the cortical pyramidal neurons (Fig. 4), application of 10 µM PAX robustly reversed the changes of single action potential waveform caused by the BK-D434G mutation and efficiently suppressed the hyperactive PCs in the BK^*DG/WT*^ mice (Fig. 6*D-H* and Table S1). Collectively, our electrophysiological characterizations demonstrated that BK-D434G GOF can also induce hyperexcitability in PCs by accelerating after-hyperpolarization and facilitating Na_V_ channel deinactivation. Sustained hyperexcitability in BK-D434G PCs may ultimately induce stress to the PCs, leading to morphological changes and contributing to the observed locomotor defects in the BK-D434G mutant mice.

## Discussion

In this study, we show that the BK-D434G knock-in mice resembles the clinical manifestations of generalized absence epilepsy observed in the BK-D434G patients (13). The BK-D434G mice exhibited spontaneous SWDs, which can be suppressed by first line anti-absence medicines and BK channel specific blocker PAX. These findings thus strongly support that BK-D434G GOF causes absence seizures in the BK-D434G channelopathy patients.

Utilizing the BK-D434G knock-in mice, we uncovered the cellular pathophysiology of the GOF BK mutation in inducing epilepsy and movement disorders. We found that BK-D434G causes hyperexcitability in both cortical pyramidal neurons and cerebellar Purkinje cells, in which BK channels are highly expressed (2, 3). BK channels are usually form protein complexes with VGCCs in the central nervous system (9). With enhanced Ca^2+^ sensitivity, BK-D434G GOF mutation will be rapidly activated following membrane depolarization and Ca^2+^ entry from the VGCCs. The enhanced BK channel activity will accelerate fAHP as evidenced by significant shortening of ADP90 (Figs. 3C, 3D, 6F, 6G) and enhanced amplitude of AHP (Figs. 3C, 3E, 6F, 6H). The accelerated fAHP in the BK-D434G neurons can facilitate the recovery of Na_V_ channels from inactivation and promote activation of HCN channels, thereby increasing membrane excitability (Fig. 3F), which collectively leads to enhanced firing and hyperexcitability (Figs. 3A-B and 6D-E).

Abnormal oscillatory rhythms within the cortico-thalamic system are generally believed to be responsible for absence seizure ictogenesis (30, 36). The hyperexcitability of BK-D434G cortical pyramidal neurons observed in this study supports the importance of cortical excitability in absence seizure pathogenesis. Future studies need to be done to comprehensively characterize the excitabilities of the different types of neurons in the cortico-thalamic system and illustrate the circuit basis of absence seizure ictogenesis in the BK-D434G mice. It is interesting to observe that GABAergic Purkinje cells from the BK-D434G mice are also hyperactive (Fig. 6D-H). One hypothesis to explain BK GOF-induced hyperexcitability is that the GOF mutations would decrease the excitability of inhibitory neurons, thereby leading to disinhibition of neuronal networks and subsequently hyperexcitability (13). Our observation that the hyperexcitability of the BK-D434G Purkinje cells suggests that the inhibitory neurons with high expression of BK GOF mutations, instead of reducing their excitability, would increase their excitability. The enhanced GABA release will augment inhibitory inputs and switch the downstream neurons in a circuit into a bursting mode, thereby causing hypersynchronization. In the future, it is therefore important to elucidate the contributions BK GOF to membrane excitability in other inhibitory neurons that have different BK channel expression levels.

In addition to having spontaneous absence seizures (Fig. 1), the BK-D434G mice are also more susceptible to PTZ-induced tonic-clonic seizures (Fig. S3). It is likely that BK-D434G GOF may also enhance the excitability of the other neurons outside of the cortico-thalamic system such as hippocampal pyramidal neurons and dentate gyrus granule cells (12). Future investigations of the excitabilities of these neurons from the BK-D434G mice will shine light on understanding the neuronal and circuit basis of developing tonic-clonic seizures in some of the refectory and/or pharmaco-resistant absence seizure patients (45).

Despite clinical applications of the first-line anti-absence medicines including ethosuximide (ESM) and valproate since 1950s (46-48), 30% of absence epilepsy patients are pharmaco-resistant and 60% of them are affected by severe neuropsychiatric comorbidities, including attentional, mood, cognitive and memory impairments (30, 49). While human genetics and animal models have shown that VGCCs and GABA_A_ receptor chloride channels contribute to the etiology of absence epilepsy (36, 50), the contributions of potassium channels to absence epilepsy pathogenesis are still elusive. In this study, we showed that PAX, a BK channel blocker, can effectively suppress BK-D434G induced hyperexcitability and absence seizures (Fig. 4), as well as PTZ-induced tonic-clonic seizures (Fig. S4). This is consistent with the previous findings that PAX can alleviate convulsant drug-induced generalized epilepsy (39, 40) and spontaneous seizures in an Angelman syndrome mouse model with enhanced BK channel activity (41). Our current study thus demonstrated that targeting BK channels could be a novel strategy to mitigate absence epilepsy. Future investigations are needed to examine if pharmacological inhibition of BK channels can be a general strategy to treat different BK GOF channelopathy and could be used to treat pharmaco-resistant absence epilepsy. Of course, better BK inhibitors also need to be developed because PAX’s anti-absence effect vanished in 30 minutes after injection due to its poor pharmacokinetics (Fig. 4*H*) (51).

The BK-D434G mice showed severe locomotor defects as examined using open field, balance beam, rotarod and gait analysis (Fig. 5), albeit no obvious sign of PNKD observed. This is different from the clinical observation in which twelve out of sixteen BK-D434G patients had PNKD (13). This discrepancy is likely due to the organism difference between mice and humans. It is also possible that the frequent absence seizures in the BK-D434G mice complicate the detection of PNKD, which is not trivial to monitor in mouse models (52). Nevertheless, the hyperexcitability and morphological changes of the BK-D434G Purkinje cells explicitly demonstrated the involvement of the cerebellum in the pathogenesis of PNKD. As BK-D434G induced hyperexcitability leads to Purkinje cell morphological changes (Fig. 6A-C), we are not clear if acute administration of PAX or the first line anti-absence medicine could mitigate the motor defects. Long-term drug treatment starting in early developmental stages is needed to examine if BK inhibition can also be used to treat movement defects. Future investigations using neuronal type-specific knockin of D434G in mice are needed to further dissect the pathophysiological mechanism of BK-D434G in PNKD and develop corresponding therapies.

Taken together, the BK-D434G knock-in mice advanced our mechanistic understanding of the pathophysiology of BK GOF in epilepsy and movement disorders. The mechanistic insights gained in this study and our attempts to use PAX to treat absence seizures will shine light on developing novel therapies to mitigate absence epilepsy and movement disorders, as well as designing precision medicine to treat BK GOF channelopathy.

## Methods

### Origin of the mouse lines used

BK-D434G mutation mice were generated by homologous recombination in embryonic stem cells and implanted in C57Bl/6J blastocysts using standard procedures. The targeting vector was designed to flank the D434G mutation with a neomycin (Neo) selection cassette with loxP sites after exon 10 of the *KCNMA1* gene (Fig. S1*A*). Chimeric mice were crossed to C57Bl/6J females (Jackson Labs). Germline transmission generated *BK*^*+*^*BK*^*D434G*^ (*BK*^*DG/WT*^) mice. Germline transmission was determined by genotyping PCR of mouse tail DNA (Fig. S1*B*), using primers pKCNMA1_genotyping F1 (5’-GTGCCTAGAGGTGGCTGGGAATTAG-3’) and pKCNMA1_genotyping R1 (5’-CCTCTCCTACGGTGGTAAAGTATCC-3’) for the wildtype allele (342 base pairs, bp) and the floxed allele (455 bp). The F1 hybrids were crossed to C57Bl/6J β-actin Cre mice to excise the Neo cassette. The D434G mutations were confirmed by primers pKCNMA1_sequencing F2 (5’-GCTGAGTGGGGAGATGTATTGCTTC-3’) and pKCNMA1_sequencing R2 (5’-ACCTAAGGAGCCAGCACCAATCAT-3’). The BK-D434G mice were then backcrossed to C57Bl/6J mice for five generations.

For all behavioral experiments, *BK*^*DG/WT*^ males were bred with *BK*^*DG/WT*^ females. Animals were housed at a constant 24 °C in a 12 h light–dark cycle (lights on at 07:00) with *ad libitum* food and water. Both males and females were used for *in vivo* and *in vitro* analysis. Mouse handling and usage were carried out in a strict compliance with protocols approved by the Institutional Animal Care and Use Committee at Duke University, in accordance with National Institutes of Health guidelines. PCR genotyping was performed using tail DNA extraction.

### Simultaneous Video-EEG Recording and Analysis

Mice with age of 1 to 6 months were anesthetized with 1∼2% isoflurane and mounted on a stereotaxic device (Kopf Instruments). A mouse electroencephalogram (EEG) headstage (#8201, Pinnacle technology Inc., Lawrence, KS, USA) was affixed to the skull with three screws, which served as differential recording leads on the frontal, parietal, and cerebellar cortex. The headstage was subsequent secured to the skull by the dental cement and the animal could recover for 5 days prior to EEG recording. EEG recordings were collected by a preamplifier with 100x gain and high pass filtered at 1.0 Hz (#8200-SE, Pinnacle technology Inc.), accompanied by spontaneous video monitoring on the top of the chamber (Logitech C920 HD Pro Webcam, 24 frames per second). For drug treatment test on the *BK*^*DG/DG*^ mice, a single dose of paxilline (0.35 mg/kg, i.p.), ethosuximide (150 mg/kg, i.p.), valproate (200 mg/kg, i.p.) or saline control was injected into the mice after 1 hour of recording as baseline. Data were acquired with an analog□to□digital converter (PCI□6221; National Instruments, Austin, TX, USA) to a desktop computer. A custom code written in MATLAB (MathWorks, Natick, MA, USA) was used to visualize the raw EEG recording trace and plot the power spectra using the Fast Fourier Transform (FFT) within the frequency range of 1-20 Hz. The numbers of SWD event were calculated using previously described methods (53).

### Video based motion analysis

The video-based motion was analyzed using a similar method previously described (54). A custom-written MATLAB code was used to analyze the video recordings of freely moving mice in the EEG recording chamber. The videos were first down-sampled to 1 frame per second, and then converted to gray 8-bit images (Fig. S2*A*, upper panel). Since the mouse is darker than the background in the gray images, we conducted the image segmentation 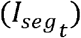 of the mouse at time *t* by

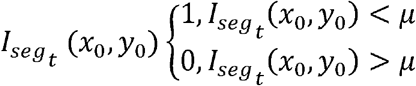

where (*x*_0_,*y*_0_) is the coordinate of the image, and *μ* is the threshold, the value of which was empirically set to be 10% of the darkest intensity (255) of the 8-bit image. A representative result of the segmentation images is shown in Fig. S2*A*, middle panel.

To get the total movement of the mice over time, we obtained a subtracted image 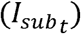 by substrating two sequential frames

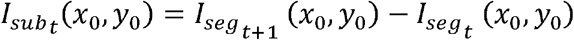

where 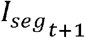 is the next sequential frame of 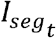. The subtracted images were shown in Fig. S2*A*, bottom panel. Pixels changed above the empirical threshold (300 pixels) were designated as a motion status.

### PTZ-induced seizure model

We performed intraperitoneal (i.p.) injection of 40 mg/kg pentylenetetrazole (PTZ; Sigma, MO, USA) and then immediately placed animals in a chamber and started video recording. PTZ-induced seizures were scored according to a modified Racine scale (55): 0, normal behavior, no abnormality; 1, immobilization, lying on belly; 2, head nodding, facial, forelimb, or hindlimb myoclonus; 3, continuous whole-body myoclonus, myoclonic jerks, tail held up stiffly; 4, rearing, tonic seizure, falling down on its side; 5, tonic-clonic seizure, falling down on its back, wildly rushing and jumping. 6: death. Score 4 and above are considered as generalized seizures. The latency to develop generalized seizure and the duration of the generalized seizure was measured based on the videos.

### Electrophysiology

For the recording performed in brain slice, acute slice preparations were as described previously (56). Briefly, *BK*^*WT/WT*^ and *BK*^*DG/WT*^ mice (postnatal day 15-24) were anesthetized with isoflurane and decapitated. For the recording in different brain regions, the section orientation is different. For the recording in the cortex, 300 µm coronal sections were prepared. For the cerebellar Purkinje cells, 250 µm sagittal slices were prepared. The brain slices were cut in ice-cold NMDG aCSF containing (in mM): 92 NMDG, 2.5 KCl, 1.2 NaH_2_PO_4_, 30 NaHCO_3_, 20 HEPES, 25 glucose, 5 sodium ascorbate, 2 thiourea, 3 sodium pyruvate, 10 MgSO_4_·7H_2_O, 0.5 CaCl_2_·2H_2_O (Titrated pH to 7.3-7.4 using concentrated HCl). The slices were then incubated in HEPES holding solution (in mM): 92 NaCl, 2.5 KCl, 1.2 NaH_2_PO_4_, 25 NaHCO_3_, 20 HEPES, 25 glucose, 5 sodium ascorbate, 2 thiourea, 3 sodium pyruvate, 2 MgSO_4_·7H_2_O, 2 CaCl_2_·2H_2_O) for 60-min at room temperature. After incubation, the slices were transferred to a recording chamber and superfused (3 mL min^-1^) with artificial cerebrospinal fluid (aCSF) at 33 □. (in mM): 124 NaCl, 2.5 KCl, 1.2 NaH_2_PO_4_, 24 NaHCO_3_, 5 HEPES, 12.5 glucose, 2 MgSO_4_·7H_2_O, 2 CaCl_2_·2H_2_O. All solutions used for electrophysiology were equilibrated with 95% O_2_/5% CO_2_. Whole-cell recordings were performed with a MultiClamp 700B amplifier and sampled at 10 kHz using a Digidata1550A A/D converter. All data acquisition and analyses were performed using the software pClamp 10.7 (Molecular Devices). For action potential recording, pipette resistance was 3-7 MΩ when filled with an intracellular solution containing the following (in mM): 125 K-gluconate, 15 KCl, 10 HEPES, 2 Mg-ATP, 0.3 Na-GTP, 10 disodium phosphocreatine, and 0.2 EGTA, adjusted to pH 7.25 with KOH. After GΩ-seal and membrane break-through, the membrane resting potential was monitored for 10 min until it is stabilized before recording of action potentials. For pharmacological experiments, 10 µM paxilline was added to extracellular aCSF. AP90% duration was defined by action potential duration of 90% repolarization. The fAHP size was measured as the difference between the spike threshold and voltage minimum after the action potential. First interspike interval was the time between the first and second action potential peaks. Input resistance (R_in_) was calculated from voltage deflections induced by rectangular hyperpolarizing current injections (0-100 pA). Membrane time constant (τ_m_) was obtained by fitting a single exponential function to these same hyperpolarizing voltage deflections. Membrane capacitance (C_m_) was calculated by dividing τ_m_ by R_in_. AP amplitude was calculated as the voltage difference between AP threshold and AP peak.

### Histology

Mice were transcardially perfused with phosphate-buffered saline (PBS) followed by 4% paraformaldehyde. The brain was removed and post-fixed in 4% paraformaldehyde overnight at 4°C and dehydrated in 30% sucrose for 48 h. Sagittal section (50 μm) containing the cerebellum Purkinje cells were collected by using a cryostat (Leica CM1900). The sections were rinsed 3 times with PBS for 10 min each and blocked with 5% goat serum and 0.3% Triton X-100 for 2 hours at room temperature and incubated for overnight at 4°C with following primary antibodies: anti-calbindin (1:1000, mouse, Sigma Aldrich, #C9848). After 3 rinses with PBS for 10 min, secondary antibodies (1:1000, conjugated with Goat anti-Mouse Alexa 594, Thermo Fisher Scientific, A-11032) were incubated for 2 hours at room temperature. Then the sections were washed 3 times with PBS for 10 min each and stained with DAPI (1:10000 of 5 mg/mL, Sigma-Aldrich). Images were acquired using a Zeiss 780 inverted confocal microscope. Representative images from at least three repeats.

### Open field test

The mice were placed in a 45 × 45 cm arena composed of four white Plexiglas walls. They could freely move in the arena for 15 min and their locomotion were continuously monitored by video recording. Locomotor activities were evaluated as the distance traveled per 5 min and the total distance by using a custom MATLAB code.

### Balance beam test

Mice were given five training trials on an 80-cm long, 7-mm small round beam elevated 30 cm above the table, as described previously (57). A video camera was placed 4-inch away from the starting point, so the hindpaws slip could be easily recorded, whereas the opposite end of the beam entered their home-cage with food pellets and bedding materials. The number of foot slips and traversal time were measured as mice traversed the beam in a test trial 24 hours after training.

### Accelerating Rotarod

The rotarod treadmills (ENV-577M, Med associates, St. Albans, VT, USA) was used to asset the motor coordination of the mice. Before testing, all mice were trained on a fixed-speed protocol at 4 rpm until they could stay on the rod for 30 s. On the same day as the training session, mice were placed on the rotarod for four-10-minute trials with 30 mins rest between trials. In each trial, the rotarod accelerated from 4 to 40 rpm at the rate of 1 rpm every 8 s, then remained at 40 rpm until the end of the trial. The time until the mouse fell from the rod was recorded as the latency to fall. The assessments were performed for four days.

### Gait Analysis

The forepaws and hindpaws of the mice were painted with non-toxic red and blue inks, respectively. After a two-minute habituation trial, each mouse could walk along a narrow, paper covered runway. The length of each stride was measured.

### Statistics

All the statistical analyses were performed using GraphPad Prism (GraphPad Software). Sample number (n) values are indicated in the results section and Figure legends. All data are presented as the mean ± standard error of the mean (s.e.m.). Sample sizes were chosen based on standards in the field as well as previous experience with phenotype comparisons. No statistical methods were used to predetermine sample size.

## Supporting information

Movie S1, BK-D434G mice have spontaneous absence seizure

Movie S2, Balance beam performance of BK-D434G mutation mice.

Movie S3, BK-D434G mutation mice exhibit motor deficiency in accelerated rotarod test

## Author contributions

H.Y. and J.C. perceived the research. H.Y. supervised the project. H.Y. and P.D. designed the experiments with critical help from J.C. and M.A.M. P.D. performed behavioral experiments and brain slice recordings. Y.Z. and P.D. conducted immunofluorescence. P.D. and Y.Z. conducted data analysis. P.D. and H.Y. wrote the manuscript.

## Acknowledgments

We are grateful to Dr. Xuechu Zhen (Soochow University, China) for providing the BK-D434G mice. We appreciate Drs. Dwight D. Koeberl and Arsen Hunanyan for their technical assistance with locomotor behavioral tests. We also thank Drs. James O. McNamara, William Wetsel, Pengfei Liang, Son Le, Trieu Le, and Zoe Shan for their critical comments on the manuscript. This work was supported by the Duke Institute for Brain Sciences (to H.Y. and M.M.) and the American Epilepsy Society Post-Doctoral Fellowship 693905 (to P.D.).

