## Supplementary Information for "Neuronal mechanism of a BK channelopathy in absence epilepsy and movement disorders"

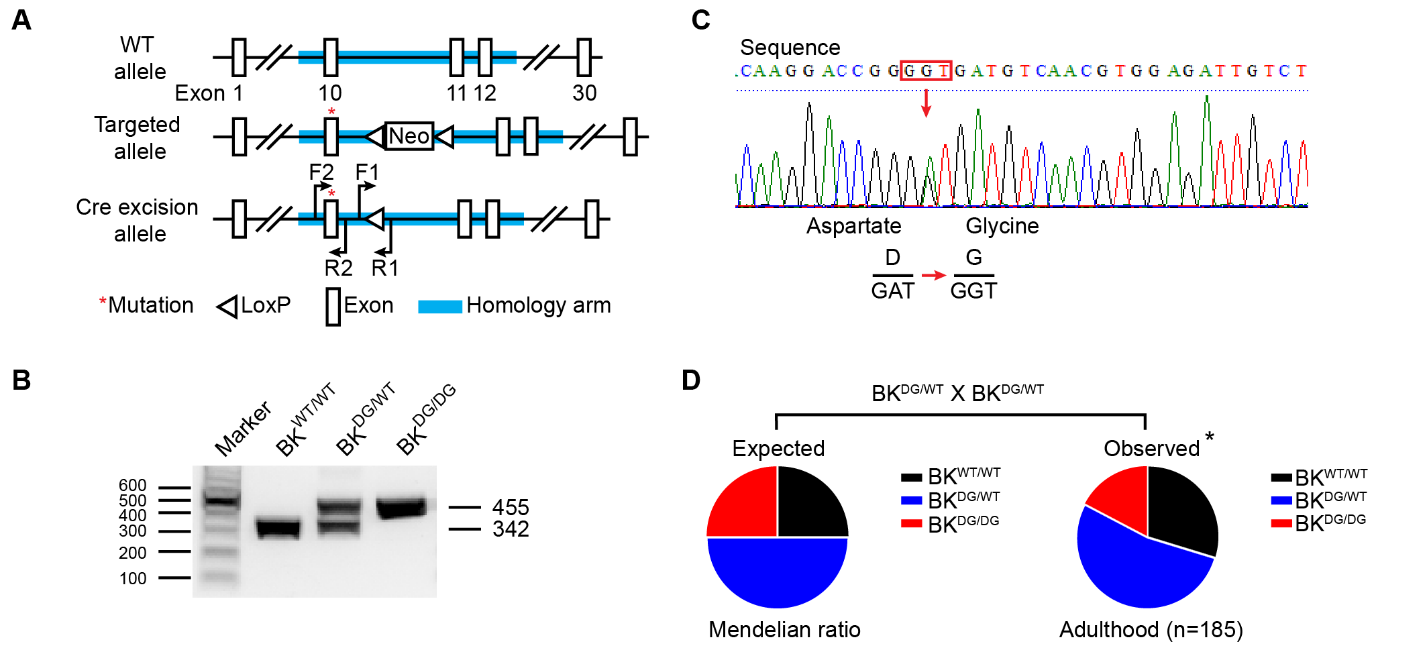
**Fig. S1.** **Generation, validation, and the survival of the BK-D434G mutant mice.** (**A**) Schematic describing the strategy to generate BK-D434G knock-in mouse model. Mice containing targeted allele were crossed to β-actin Cre mice to remove the Neo cassette. The red asterisk indicates the D434G mutation site. F1, R1 are the primers used for the PCR genotyping in panel B. F2, R2 are the primers used for sequencing confirmation in panel C. (**B**) PCR genotyping of *BK^WT/WT^*, *BK^DG/WT^* and *BK^DG/DG^* mice. (**C**) DNA sequencing of a heterozygous *BK^DG/WT^* mouse confirmed the *GAT* to *GGT* mutation in the *Kcnma1* gene, which results in the replacement of aspartate 434 with a glycine (D434G). (**D**) Reduction of homozygous BK-D434G mice from the expected Mendelian inheritance. **P* < 0.05 with the Chi-squared test.

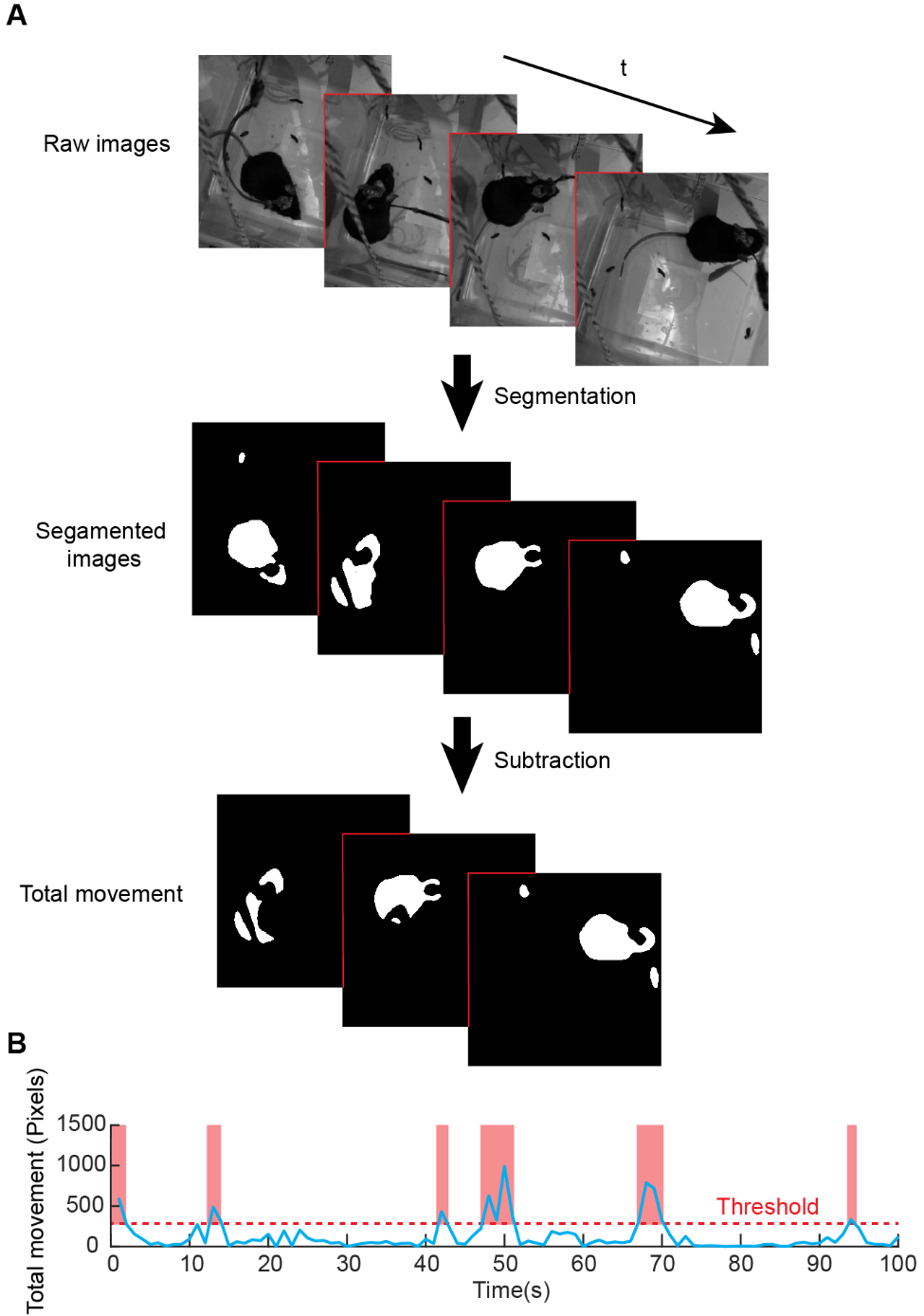
**Fig. S2.** **Automatic tracing and quantification of the total movement of freely moving mice based on video analysis****.** (**A**) Example video raw frames and automated segmentation the mice from the background using a custom MATLAB program based on the threshold. To obtain the total movement, sequential frames are subtracted to compare the difference between each frame. Note: The snapshots shown in the figure are video frames at 10 s interval for visualization purpose; for real data analysis, the interval was set to 1 s in panel **B** and the rest of the paper. See Methods for details. (**B**) Quantification of the mice total movement based on threshold-based segmentation in an example recording trial. See Methods for details on setting the threshold and data analysis.

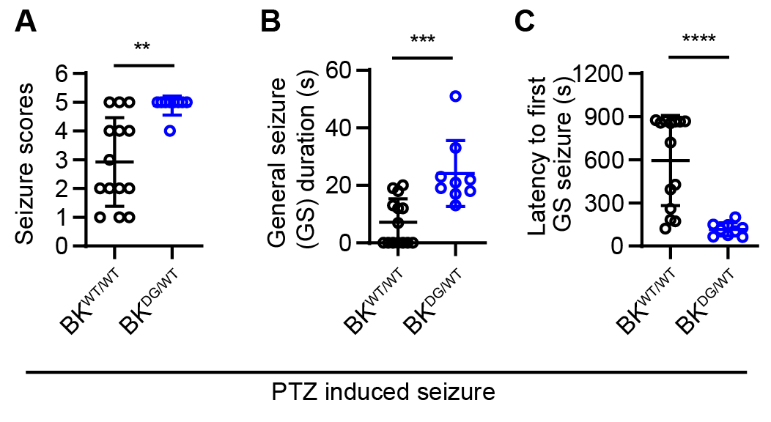

**Fig. S3. BK-D434G knock-in mice are susceptible to pentylenetetrazole (PTZ)-induced seizure.** (**A**-**C**) *BK^DG/WT^* mice were more vulnerable to 40 mg/kg PTZ-induced seizure model. *BK^DG/WT^* mice showed increased seizure score (**A**), prolonged generalized seizure (GS) duration (**B**) and reduced latency to GS (**C**). Two-tailed unpaired Student’s *t*-test: ** *P* < 0.01, *** *P* < 0.001, **** *P* < 0.0001. *BK^WT/WT^*, n = 14 mice; *BK^DG/WT^*, n = 9 mice. In all plots and statistical tests, summary graphs show mean ± s.e.m.

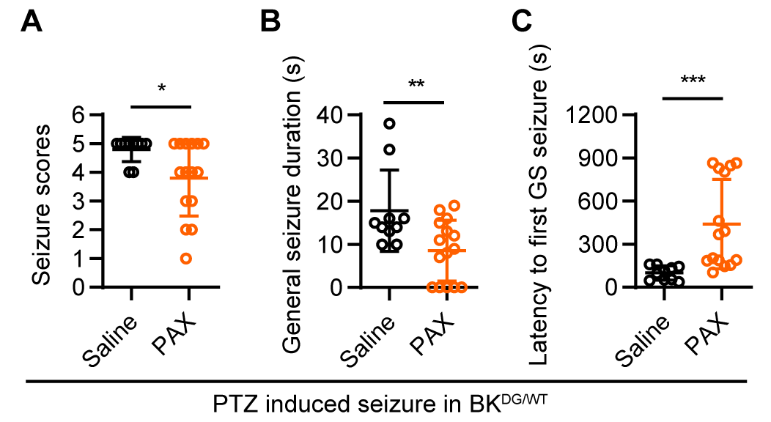
Fig. S4. Paxilline (PAX), a potent BK channel inhibitor can decrease PTZ induced seizure severity in BK-D434G mice. (A-C) Co-injection of paxilline in the heterozygous BK-D434G mice can decrease PTZ induced seizure stage score (A); shorten generalized seizure (GS) duration (B); and increase the latency to GS (C). Two-tailed unpaired Student’s t-test: * p<0.05, ** p<0.01, ***p<0.001. Saline, n = 10 mice, Paxilline, n = 15 mice. In all plots and statistical tests, summary graphs show mean ± s.e.m..

**Table S1. The membrane properties wild-type and BK-D434G neurons.**

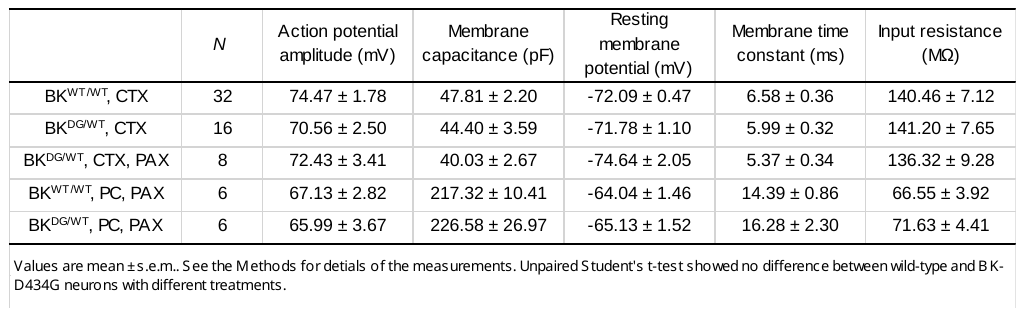

Abbreviation: cortex, CTX; Purkinje cell, PC; paxilline, PAX.

**Captions for Movies**

**Movie S1. Spontaneous absence seizure with behavioral arrest of a *BK ^DG/DG^* mouse.** Video shows a representative simultaneous video-EEG recording of a freely moving *BK^DG/DG^* mouse. The mouse had behavior arrest when the SWDs appeared and the SWDs resolved when the animal moved. Video was shown at 6 times speed.

**Movie S2. BK-D434G mutation mice had defects to traverse balance beam**

Videos show BK^WT/WT^ control (left), BK^DG/WT^ (middle) and BK^DG/DG^ (right) mice traversed through the balance beam. The BK^DG/WT^ and BK^DG/DG^ mice made more hind-limb slips and took longer time to cross the balance beam. Note that the tails of the BK^DG/WT^ and BK^DG/DG^ mice frequently wrapped around the beam to keep their balance when their hind-limbs slipped. The video is shown at actual speed.

**Movie S3. BK-D434G mice failed on accelerated rotarod**

Video shows two BK^WT/WT^ control mice, two BK^DG/WT^ mice and one BK^DG/DG^ mouse running on the accelerated rotarod. Video is shown at 5 times speed.
